# An 1,4-α-glucosyltransferase defines a new maltodextrin catabolism scheme in *Lactobacillus acidophilus*

**DOI:** 10.1101/2020.03.17.996314

**Authors:** Susan Andersen, Marie S. Møller, Jens-Christian N. Poulsen, Michael J. Pichler, Birte Svensson, Yong Jun Goh, Leila Lo Leggio, Maher Abou Hachem

## Abstract

The maltooligosaccharide (MOS) utilization locus in *Lactobacillus acidophilus* NCFM, a model for human small-intestine lactobacilli, encodes a family 13 subfamily 31 glycoside hydrolase (GH13_31), annotated as an 1,6-α-glucosidase. Here, we reveal that this enzyme (*La*GH13_31B) is an 1,4-α-glucosyltransferase that disproportionates MOS with preference for maltotriose. *La*GH13_31B acts in concert with a maltogenic α-amylase that efficiently releases maltose from MOS larger than maltotriose. Collectively, these two enzymes promote efficient conversion of preferentially odd-numbered MOS to maltose that is phosphorolysed by a maltose phosphorylase, encoded by the same locus. Structural analyses revealed the presence of a flexible elongated loop, which is unique for *La*GH13_31B and its close homologues. The identified loop insertion harbours a conserved aromatic residue that modulates the activity and substrate affinity of the enzyme, thereby offering a functional signature of this previously undescribed clade, which segregates from described activities such as 1,6-α-glucosidases and sucrose isomerases within GH13_31. Sequence analyses revealed that the *La*GH13_31B gene is conserved in the MOS utilization loci of lactobacilli, including acidophilus cluster members that dominate the human small intestine.

**IMPORTANCE:** The degradation of starch in the small intestine generates short linear and branched α-glucans. The latter are poorly digestible by humans, rendering them available to the gut microbiota *e.g*. lactobacilli adapted to the human small intestine and considered as beneficial to health. This study unveils a previously unknown scheme of maltooligosaccharide (MOS) catabolism, via the concerted action of activity together with a classical hydrolase and a phosphorylase. The intriguing involvement of a glucosyltransferase is likely to allow fine-tuning the regulation of MOS catabolism for optimal harnessing of this key metabolic resource in the human small intestine. The study extends the suite of specificities that have been identified in GH13_31 and highlights amino acid signatures underpinning the evolution of 1,4-α-glucosyl transferases that have been recruited in the MOS catabolism pathway in lactobacilli.

## INTRODUCTION

Humans are co-evolved with a diverse and vast bacterial community (1) termed the human gut microbiota (HGM), which exerts considerable effects on our health as well as metabolic and immune homeostasis (2). The gut microbiota confers metabolic activities that are not encoded by the human genome, *e.g*. the bioconversion of xenobiotics (3) and harvesting of energy from non-digestible glycans (4). The most prevalent and abundant gut bacterial phyla in healthy adults are Firmicutes, Bacteroidetes and Actinobacteria (5, 6). Complex chemical and physical gradients provide diverse ecological and metabolic niches along the gastrointestinal tract, thereby defining a biogeography for different HGM taxa. Thus, the small intestine is enriched with bacteria from the Gram-positive Lactobacillaceae family (6), especially *Lactobacillus* spp. that are ascribed health promoting properties (7, 8).

*Lactobacillus acidophilus* NCFM is one of the best characterized models for acidophilus cluster human gut lactobacilli (7, 9). This strain has been used commercially as a probiotic and synbiotic, owing to its suggested potential in improving intestinal transit during constipation as well as mucosal functions in healthy elderly (10, 11). The adaptation of *L. acidophilus* NCFM to the small intestine, which is rich in dietary carbohydrates, is highlighted by the numerous carbohydrate specific transporters and intracellular carbohydrate active enzymes (CAZymes, http://www.cazy.org/; (12)) encoded by this bacterium. The genetic and molecular basis for the high saccharolytic capacity of *L. acidophilus* NCFM has been explored for a range of non-digestible di- and oligosaccharides (13, 14) as well as plant glycosides (15).

Starch from cereals, tubers, roots and rhizomes dominates human caloric intake (16). Digestion of starch by human salivary and pancreatic α-amylases results in significant amounts of malto-oligosaccharides (1,4-α-gluco-oligosaccharides; MOS) and 1,6-α-branched limit dextrins, the latter being recalcitrant to digestion by human digestive enzymes. Therefore, these oligosaccharides represent an abundant metabolic resource for bacteria that colonize the small intestine. Starch metabolism is relatively well-studied in colonic microbiota members, e.g. from Bacteroides and Clostridia (17, 18). By contrast, α-glucan metabolism by lactobacilli that dominate the small intestine has received less attention.

Recently, we have shown that the utilization of short branched α-glucans from starch degradation by *L. acidophilus* NCFM is conferred by a cell-attached 1,6-α-debranching enzyme (*La*Pul13_14) (19). The MOS products from this enzyme or from human starch breakdown are likely internalized by an ATP-binding cassette transporter, which is conserved in MOS utilization gene clusters in lactobacilli together with amylolytic and catabolic enzymes (gene locus tags LBA1864−LBA1874 in *L. acidophilus* NCFM) (Fig. 1) (20).

**FIG 1.**
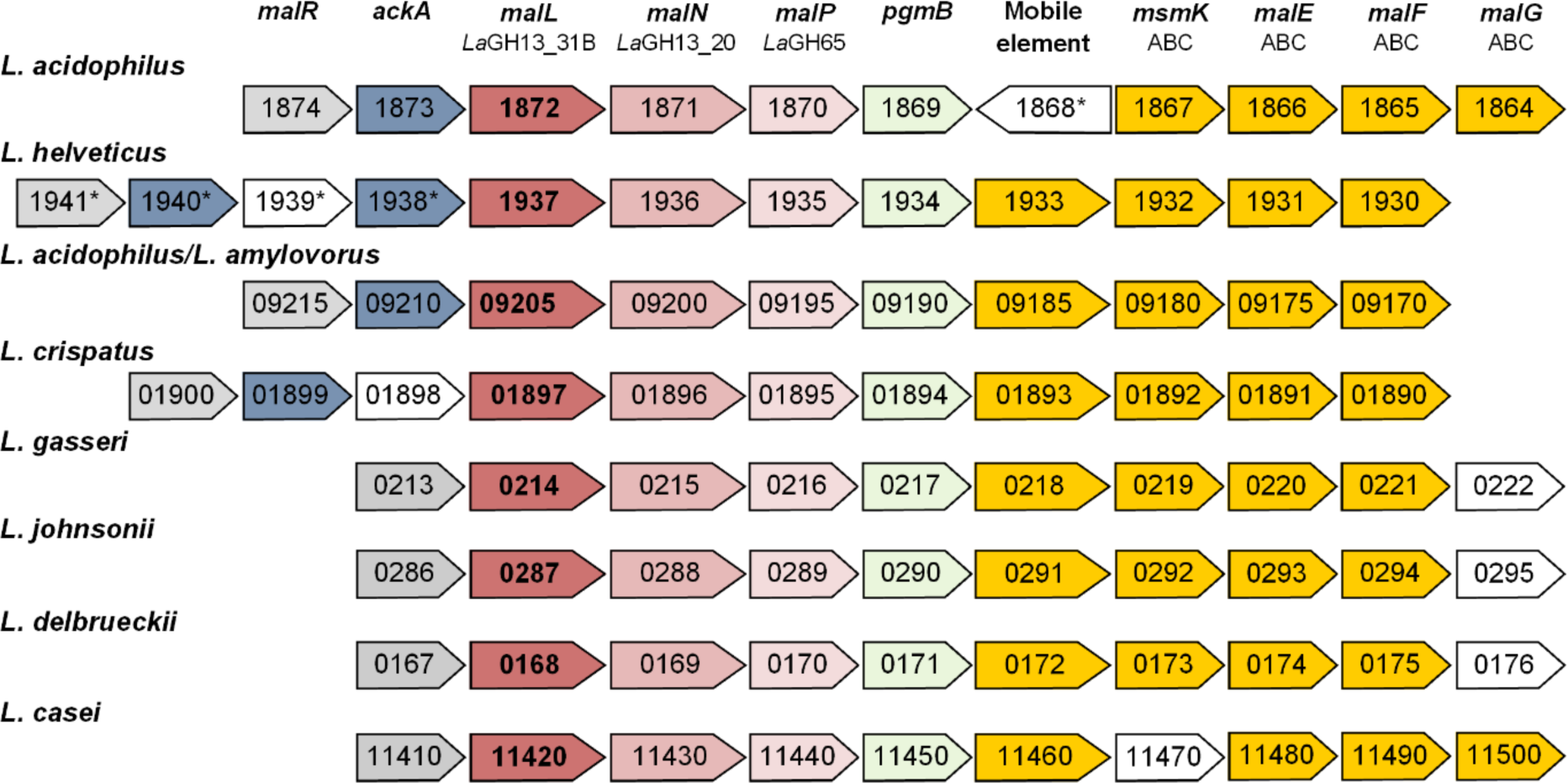
Organization of malto-oligosaccharide utilization loci in gut gut lactobacilli strains. The numbers indicated genes locus tags of the first strain mentioned in the following clusters: *<u>L. acidophilus</u>*; *L. acidophilus* NCFM, *L. acidophilus* La-14. *<u>L. helveticus</u>*; *L. helveticus* H10, *L. helveticus* CNRZ32, *L. helveticus* R0052, *L. helveticus* DPC 4571. *<u>L. amylovorus/L. acidophilus</u>*; *L. acidophilus* 30SC, *L. amylovorus* GLR1118, *L. amylovorus* GLR1112. *<u>L. crispatus</u>*; *L. crispatus* ST1. *<u>L. gasseri/L. johnsonii</u>*; *L. johnsonii* FI9785, *L. gasseri* ATCC 33323, *L. johnsonii* N6.2, *L. johnsonii* DPC6026, *L. johnsonii* NCC533. *<u>L. delbrueckii</u>*; *L. delbrueckii* subsp. *bulgaricus, L. delbrueckii* subsp. *bulgaricus* ND02. *<u>L. casei</u>*; *L. casei* BL23, *L. casei* LOCK91, *L. casei* Zhang, *L. casei* BD-II, *L. casei* ATTCC 334, *L. casei* LCZW. Transcriptional regulators (light grey), acetate kinases (blue), *La*GH13_31B orthologues (bold and red), *La*GH13_20 orthologues (pink), *La*GH65 orthologues (light pink), β-phosphoglucomutase (green), components of an ATP-binding cassette transporter (orange). The following genes are shown in white: transposase (1868 and 1939), hypothetical protein (01898), α-1,6-glucosidase (11470, 0295 and 0176). *; not present in all published genomes.

The activity of the glycoside hydrolase family 65 maltose phosphorylase (*La*GH65, LBA1870; EC 2.4.1.8) encoded by the MOS locus has been demonstrated (20). Another conserved gene within this locus (LBA1872, Fig. 1) is currently annotated as an oligo-1,6-α-glucosidase belonging to GH13 subfamily 31 (GH13_31). We have previously described another functional GH13_31 glucan-1,6-α-glucosidase from *L. acidophilus* (*La*GH13_31A) (21), which raised questions regarding the function of organization of the LBA1872 gene within the MOS utilization locus of *L. acidophilus*.

In this study, we demonstrate that LBA1872 from *L. acidophilus* NCFM encodes a MOS disproportionating enzyme (1,4-α-glucosyltransferase, henceforth *La*GH13_31B) with preference for maltotriose (M3) as an acceptor. The general reaction scheme of this disproportionating enzyme is to use a MOS with *n* glucosyl units both as a donor and an acceptor yielding two products having *n*-1 and *n*+1 units. Sequence and structural analyses have revealed a functional signature of this activity involving a dynamic loop that modulates the activity and substrate affinity of the enzyme. We expressed and kinetically characterized the two remaining MOS active enzymes (*La*GH65/LBA1870 and *La*GH13_20/LBA1871) in the MOS utilization locus of in *L. acidophilus* NCFM and showed that *La*GH13_31B acts in concert with these enzymes to enable efficient breakdown of MOS.

## RESULTS

### The MOS utilization gene cluster in *L. acidophilus* NCFM encodes a disproportionating enzyme

*La*GH13_31B was not active towards panose and isomaltose, which are diagnostic of α-(1,6)-glucosidase activity (Fig. S1). By contrast, *La*GH13_31B disproportionated MOS with a degree of polymerization (DP) of 2−7 (M2*–*M7) by one glucosyl moiety (Fig. S2A and B) and no hydrolysis products were observed. The enzyme had similar equilibrium activity profiles on M3 through M7, while M2 was a poor substrate (Fig. S2B). Moreover, the presence of amylose (DP17) or glycogen did not alter the activity profiles (Fig. S2C), establishing the specificity of *La*GH13_31B as oligosaccharide-specific 1,4-α-glucosyltransferase. Coupled enzyme assays and HPAEC-PAD were used to measure the apparent kinetic parameters of disproportionation by *La*GH13_31B, using M2*–*M4 as substrates (Table 1, Fig. S3). The best substrate (coupled enzymatic assay) was M3 owing to a higher *k*_cat_ than for both M2 and M4. Similar *K*_m_ and catalytic efficiency (*k*_cat_/*K*_m_) values on M4 were found by the coupled enzymatic assay and HPAEC-PAD analysis (Table 1). Roughly a 3-fold higher *k*_cat_/*K*_m_ was obtained on M3 as substrate using the coupled enzyme assay as compared to the HPAEC assay (Table 1), suggesting that the production of M2 in the reaction may affect the reaction kinetics. Notably, the catalytic efficiency (*k*_cat_/*K*_m_) on M2 was about 4,700-fold lower compared to M3 owing to a very high *K*_m_ for M2 (Table 1).

**TABLE 1.**
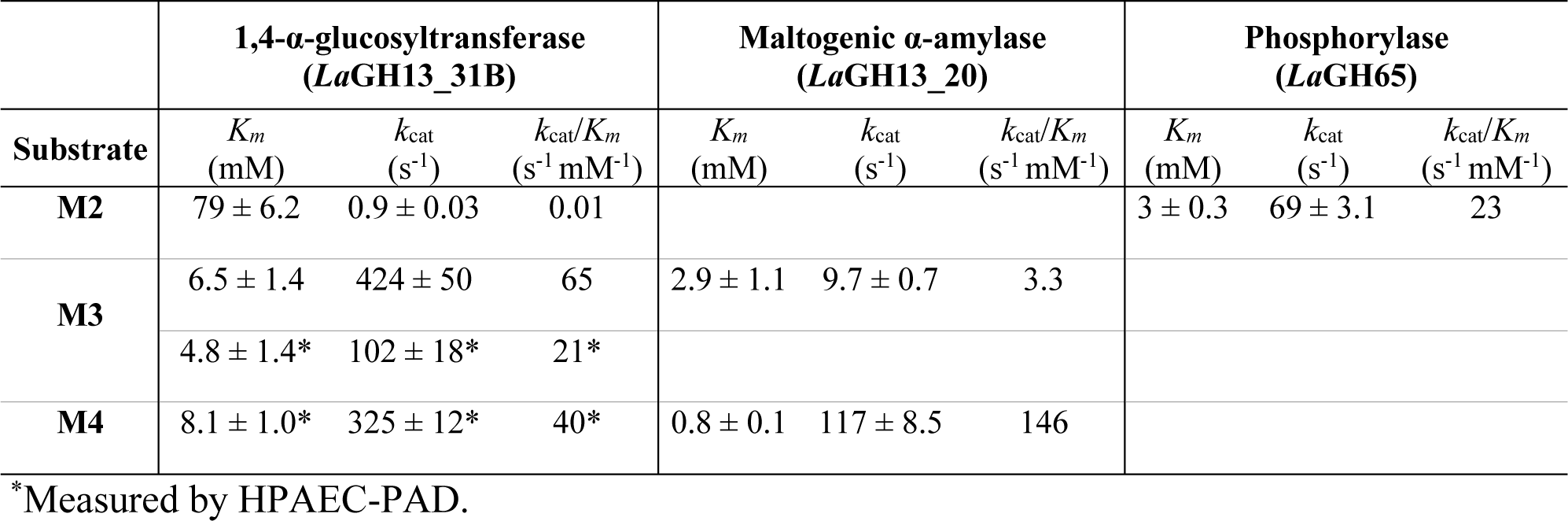
Kinetic parameters of enzymes in the maltodextrin utilization gene cluster in *L. acidophilus* NCFM. See Fig. S3 for Michaelis-Menten plots and Fig. 5 for the reactions catalyzed by these enzymes.

|  | **1,4-α-glucosyltransferase**<br>**(LaGH13_31B)** |  |  | **Maltogenic α-amylase**<br>**(LaGH13_20)** |  |  | **Phosphorylase**<br>**(LaGH65)** |  |  |
| --- | --- | --- | --- | --- | --- | --- | --- | --- | --- |
| **Substrate** | $K_m$<br>(mM) | $k_{cat}$<br>(s<sup>-1</sup>) | $k_{cat}/K_m$<br>(s<sup>-1</sup> mM<sup>-1</sup>) | $K_m$<br>(mM) | $k_{cat}$<br>(s<sup>-1</sup>) | $k_{cat}/K_m$<br>(s<sup>-1</sup> mM<sup>-1</sup>) | $K_m$<br>(mM) | $k_{cat}$<br>(s<sup>-1</sup>) | $k_{cat}/K_m$<br>(s<sup>-1</sup> mM<sup>-1</sup>) |
| **M2** | 79 ± 6.2 | 0.9 ± 0.03 | 0.01 |  |  |  | 3 ± 0.3 | 69 ± 3.1 | 23 |
| **M3** | 6.5 ± 1.4 | 424 ± 50 | 65 | 2.9 ± 1.1 | 9.7 ± 0.7 | 3.3 |  |  |  |
|  | 4.8 ± 1.4* | 102 ± 18* | 21* |  |  |  |  |  |  |
| **M4** | 8.1 ± 1.0* | 325 ± 12* | 40* | 0.8 ± 0.1 | 117 ± 8.5 | 146 |  |  |  |

\*Measured by HPAEC-PAD.

**TABLE 2.**
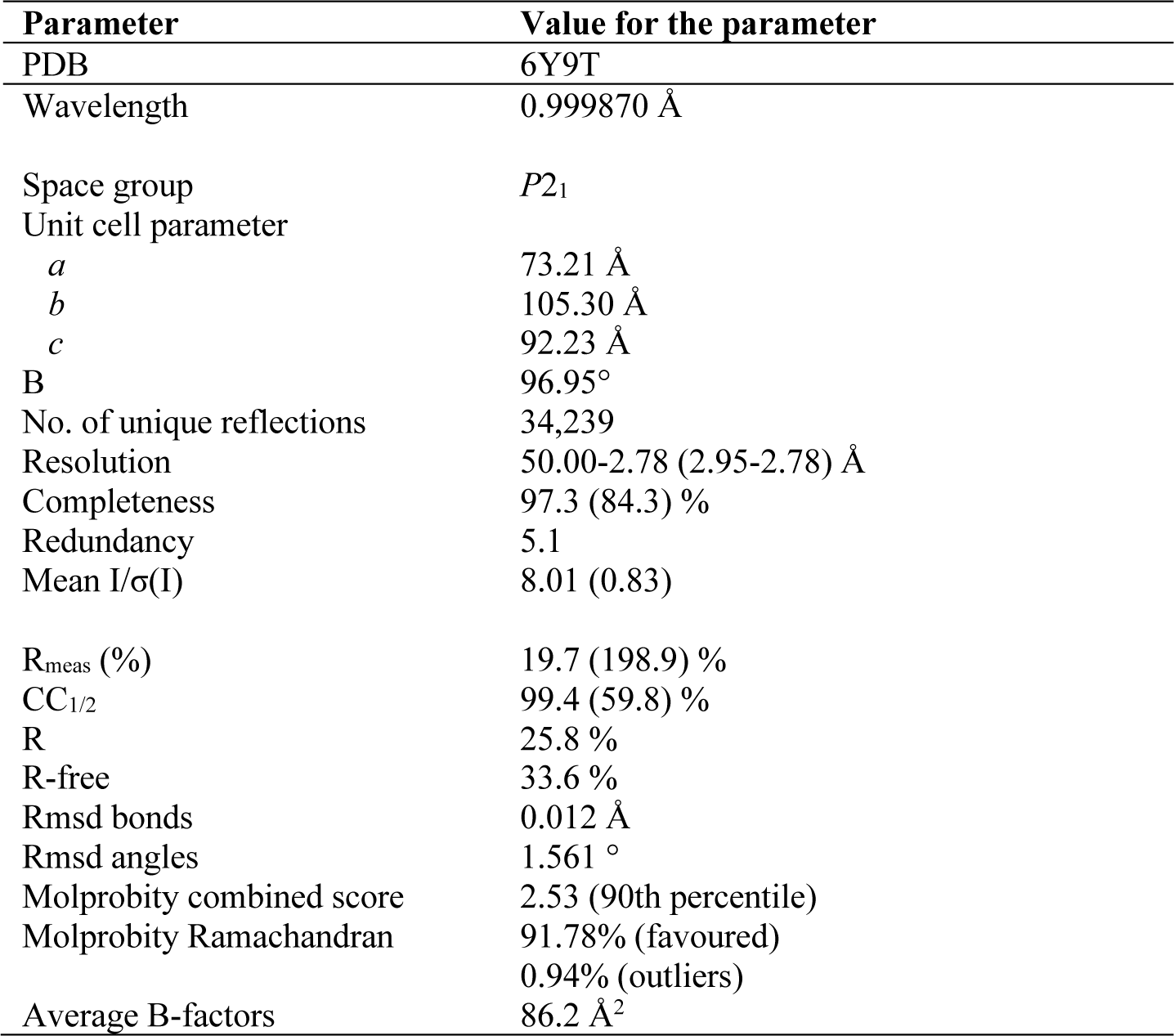
Crystallographic data collection and refinement statistics.

### Enzymology of MOS degradation in *L. acidophilus* NCFM

To dissect the role of *La*GH13_31B in MOS metabolism, we carried out kinetic analyses on the maltose phosphorylase *La*GH65 and the maltogenic α-amylase *La*GH13_20, both encoded by the same MOS locus as *La*GH13_31B (Fig. 1). The efficiency of *La*GH65 on M2 was 1,500 fold higher than of *La*GH13_31B (Table 1). Comparing the efficiency of *La*GH13_20 and *La*GH13_31B on M3 and M4 revealed that these enzymes had a reciprocal preference to these MOS. While M3 was a fairly poor substrate for *La*GH13_20, owing to a low *k*_cat_, the efficiency of *La*GH13_31B towards this substrate was 20-fold higher (coupled assay) due to a 44-fold higher *k*_cat_ (Table 1). Conversely, M4 was the preferred substrate for *La*GH13_20 as compared to *La*GH13_31B, owing mainly to a 10-fold lower *K*_m_. Finally, the action on MOS by *La*GH13_31B in the presence of both *La*GH65 and *La*GH13_20 was tested using HPAEC-PAD analysis (Fig. S4). The mixture, resulted in effect break down and the accumulation of maltose and some glucose, which was not observed for *La*GH13_31B alone.

### *La*GH13_31B displays a more open active site with a potentially dynamic loop compared with GH13_31 hydrolases

The structure of *La*GH13_31B is the first to represent the 1,4-*α*-glucosyltransferase activity within GH13_31. *La*GH13_31B shares the catalytic machinery and the overall domain structure of amylolytic GH13 enzymes, namely a (β/α)_8_ catalytic domain A (residues 1–477; catalytic residues D198, E255, and D334) with two inserted domains (domain B, residues 100–169; domain B′, residues 373–459), and a domain C composed primarily of β-sheets (478–550) (Fig. 2A). A Ca^2+^ binding site formed by side chains of D20, N22, D24, D28 and main chain carbonyls of I26 and H73, is identified in a similar location to counterparts present in some GH13_31 structures (22).

**FIG 2.**
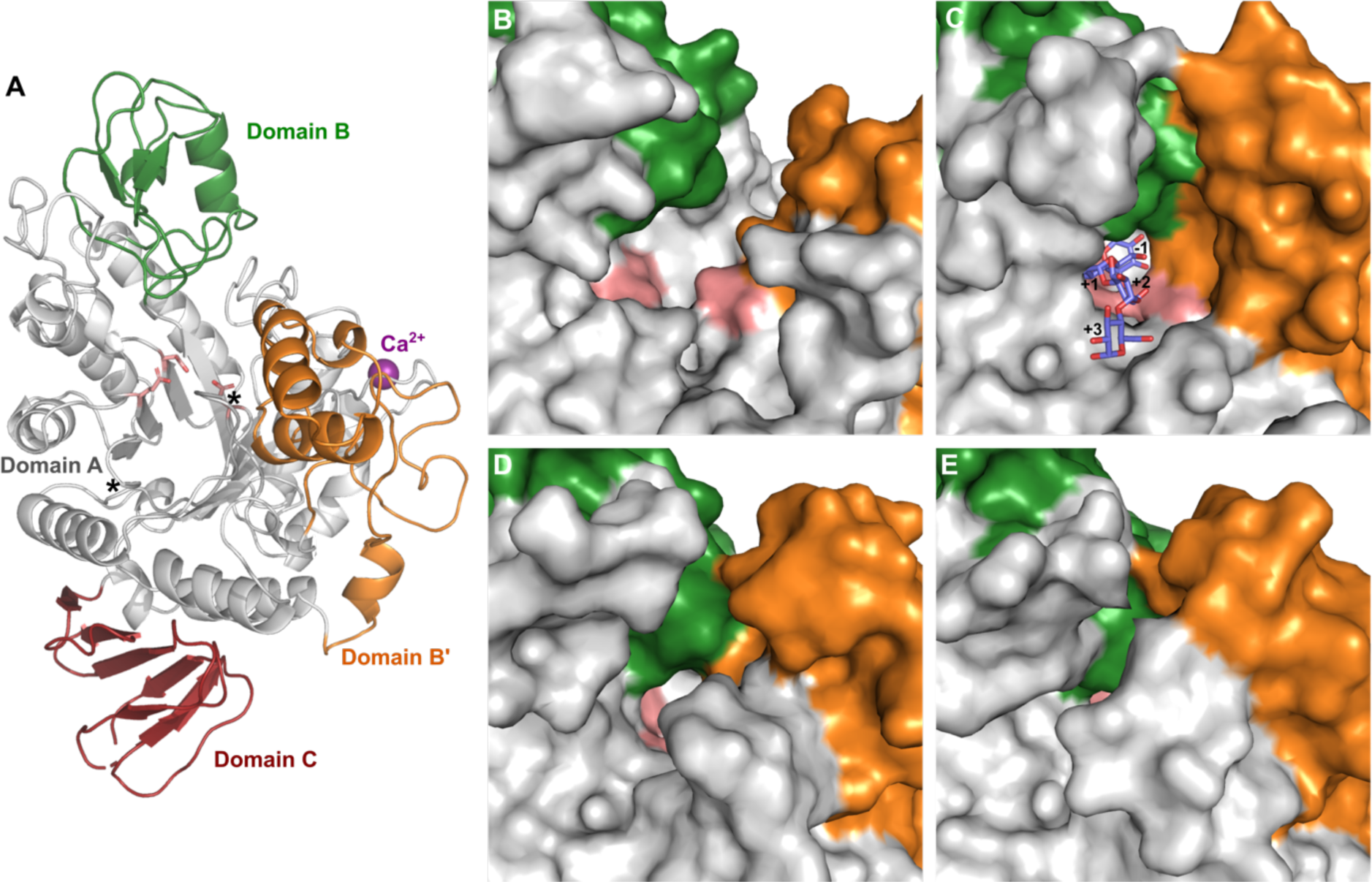
(A) Overall structure of *La*GH13_31B. Domain A (residues 3–100, 170–373, and 458– 477, grey), domain B (residues 101–169, green), domain B’ (residues 373–457, in orange) and domain C (residues 478–550, red). The catalytic residues (nucleophile, D198; general acid/base, E255; transition stabilizer, D334) are in light red, and Ca^2+^ as a purple sphere. The loop residues 286–295 (marked with asterix) were not solved. (B–E) Comparison of the orientation of the B′-domain of selected GH13_31 structures with the same domain colour as in (A): (B) *La*GH13_31B (PDB: 6Y9T), (C) *Bsp*AG13_31A (PDB: 5ZCE) with M4 as blue sticks and subsites labelled, (D) oligo-1,6-*α*-glucosidase from *Bacillus cereus* (PDB: 1UOK), (E) *α*-glucosidase (sucrase-isomaltase-maltase) from *Bacillus subtilis* (PDB: 4M56).

A DALI search identified a GH13 subfamily 29 trehalose-6-phosphate hydrolase (PDB: 5BRQ) as the closest structural homologue to *La*GH13_31B closely followed by GH13_31 enzymes, all sharing 32–36% sequence identity (Table S1). Based on a phylogenetic analysis of GH13_31 sequenence from lactobacilli and characterized GH13_31 members (Fig. 3A), the recently characterised α-glucosidase from *Bacillus* sp. *Bsp*AG13_31A (22) and a *Geobacillus* α-glucosidase (23) were found to be the closest structurally characterized enzymes.

**FIG 3.**
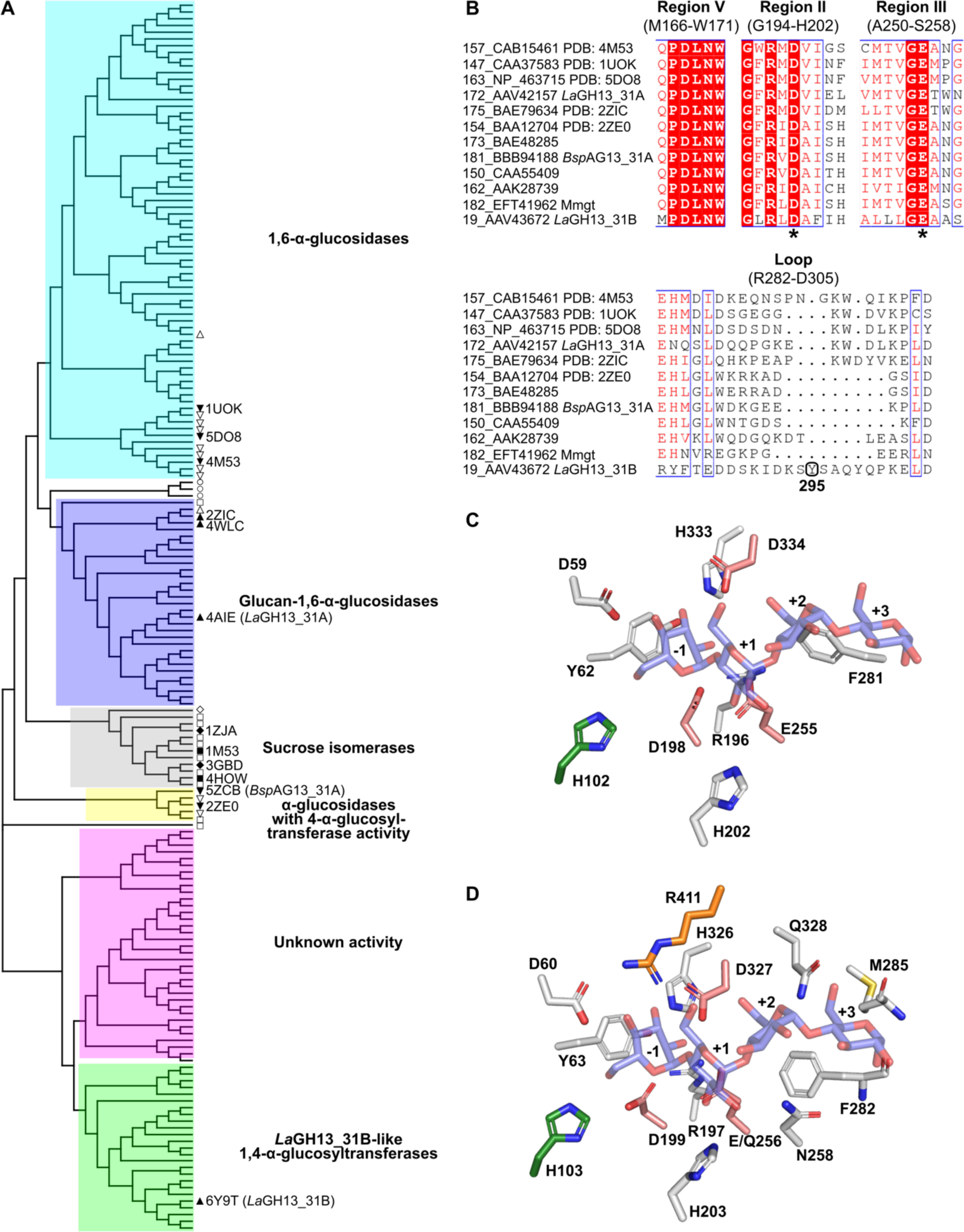
(A) Phylogenetic tree of GH13_31 protein sequences from lactobacilli and GH13_31 enzymes annotated as characterized in the CAZy database. The phylogenetic tree is based on a structure-based multiple sequence alignment of 183 sequences (Table S2, Fig. S6). Characterised GH13_31 sequences from CAZy are labelled according to the taxonomic order of the organism they originates from (Δ, Lactobacillales; ▽, Bacillales; ○, Bifidobacteriales; □, Enterobacterales; ◊, Other Gammaproteobacteria) and solid labels together with a PDB entry denote structurally characterized members (accessions and source organisms are in Table S2). (B) Excerpts of conserved GH13 regions II, III (48), and V that offers a signature discriminating oligo-1,6-α-glucosidases and neopullulanases (24), including selected characterised sequences (sequence numbers 173, 181, 150, 162 represents the *Bsp*AG13_31A-like clade) (see Fig. S6 for full alignment). The catalytic nucleophile and the general acid/base are indicated by asterisks. An excerpt of the Y295A-harbouring loop of *La*GH13_31B is shown. (C) and (D) comparison of *La*GH13_31B (M4 from PDB: 5ZCE superimposed; blue stick representation) and *Bsp*AG13_31A (PDB: 5ZCE) active site residues, respectively. The colouring is as in Fig. 2, with the catalytic residues shown in light red.

A structural difference from other GH13_31 structures is the positioning of the domain B′, which is tilted away from the catalytic domain to create a more open cleft-like active site in *La*GH13_31B. By contrast, the rest of the structures from hydrolases within GH13_31 have the domain B′ packing closely to the catalytic domain resulting in a pocket-shaped active site architecture (Fig. 2B–E).

Electron density was lacking for a long loop at the entrance to the active site, hence residues 286–295 are not solved in the structure (Fig. S5). Notably, the corresponding but shorter loop in *Bsp*AG13_31A was suggested to be flexible in *Bsp*AG13_31A based on the the poor electron density of the loop and ligand-binding dependent conformational changes in this loop (22).

### Sequence alignment and phylogenetic analysis

To map the taxonomic distribution of *La*GH13_31B-like sequences and to identify possible functional signatures of the transferase activity, we performed a sequence and phylogenetic analysis. *La*GH13_31B and close homologues segregate in a distinct clade populated by *Lactobacillus* sequences (Fig. 3A) and the genes encoding these sequences are exclusively organized similary to *La*GH13_31B in MOS utilization loci (Fig. 1). Sequences populating the *La*GH13_31B clade lack the signature of 1,6-α-glucosidases within region II of GH13 enzymes, i.e. a valine following the catalytic nucleophile, and they possess a neopullulanase (α-1,4-hydrolase) motif instead of a 1,6-α-glucosidase motif in region V, indicative of α-1,4-linkage activity (24) (Fig. 3B). Another distinguishing feature is the occurence of small amino acid residues following the catalytic general acid/base in the conserved region III (Fig. 3B), which results in a more open entrance to the active site as compared to 1,6-*α*-glucosidases. Furthermore, this difference at region III is reflected by differences in the amino acid residues that define subsites +2 and +3, when the structure of *La*GH13_31B is compared with *Bsp*AG13_31A (PDB: 5ZCE) (Fig. 3C and 3D). Thus, less interactions to substrate will be formed at the +2 and +3 subsites or a different mode of substrate binding maybe possible in *La*GH13_31B.

Another interesting feature of the *La*GH13_31B clade, is an insertion in a loop region (R282-D305) that harbors a conserved aromatic residue (Y295, *La*GH13_31B numbering), which is absent in the sequences of the other enzyme specificities of GH13_31 (Fig. 3B, Fig. S6). Interestingly, this loop region is the same region, which is disordered in the crystal structure described above. This loop could provide interaction with substrate at subsites +2 and +3, hence compensating for the fewer substrates interactions observed for *La*GH13_31B in comparison with *Bsp*AG13_31A (Fig. 3C and 3D).

### The unique extended loop of *La*GH13_31B contributes to substrate binding affinity

In order to examine the functional role of the distinctive loop insertion and the conserved Y295 in *La*GH13_31B, we mutated this residue to an alanine. The mutant enzyme had similar unfolding profile as the wildtype enzyme, precluding a gross change in the overall folding and stability (Fig. 4A). The activity on M3 was greatly reduced and no disproportionation products larger than M4 were observed in the TLC analysis (Fig. 4B). The loss of activity was confirmed by specific activity measurements and was more severe for larger substrates based on 40-fold and 140-fold loss of activity on M2 and M3, respectively, using the coupled assay (Fig. 4C). Moreover, the change in kinetic signature suggests a substantial loss in substrate affinity (Fig. 4D). These data support an important role of the flexible loop and the conserved aromatic residue it harbors in substrate binding. This loop may provide favourable binding interactions to define a dominant +2 subsite that governs anchoring the substrate at this site, which would favour unproductive binding of M2 at subsites +1 and +2. This is consistent with the high *K*_m_ on M2 and the excellent affinity to M3 as well as the larger reduction of activity toward M3 than M2 for the *La*GH13_31B Y295A mutant (Fig. 4C).

**FIG 4.**
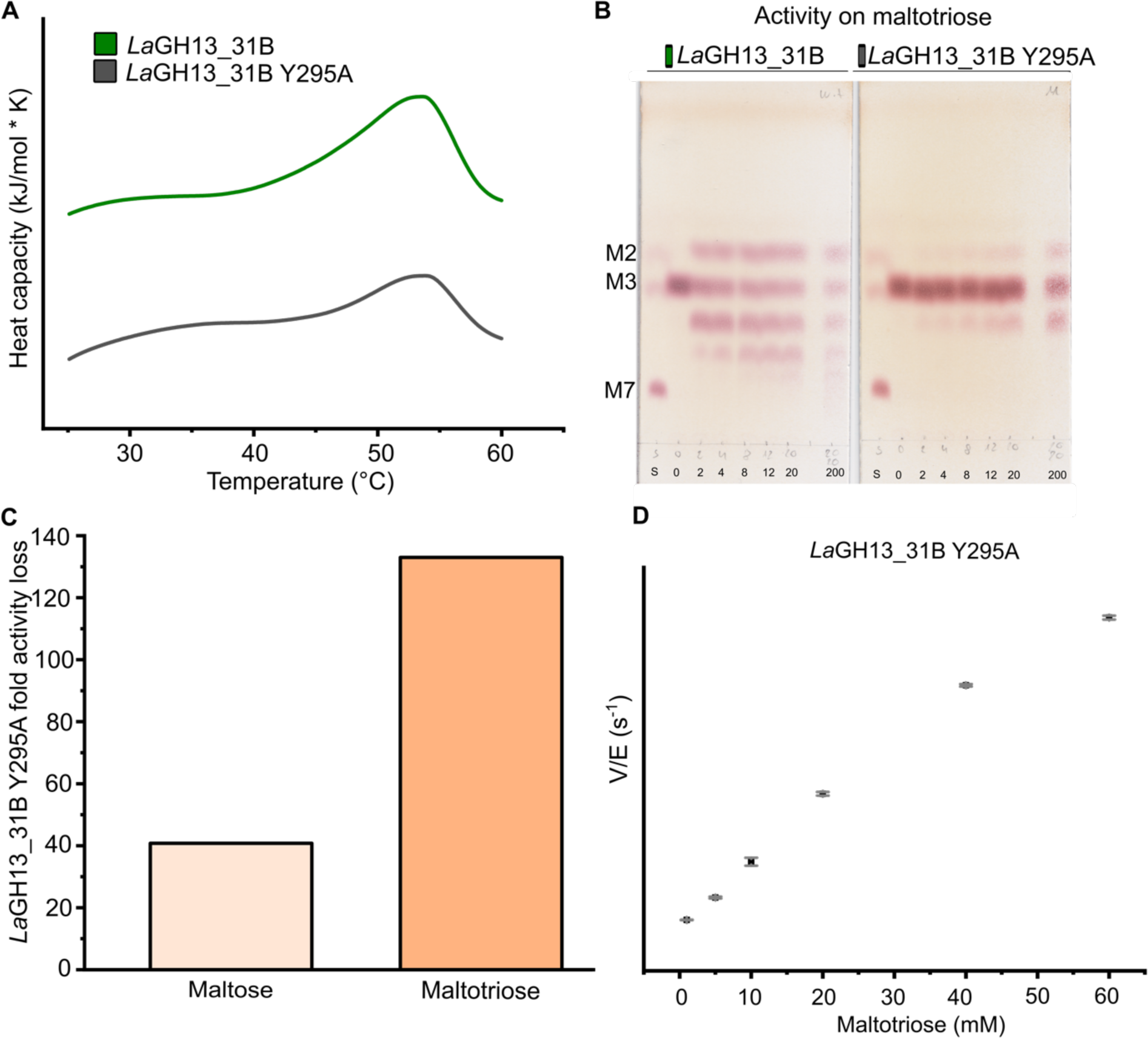
Thermal stability and activity of *La*GH13_31B and the mutant *La*GH13_31B Y295A. (A) Reference and baseline subtracted DSC thermograms showing similar unfolding and comparable thermal stabilities of *La*GH13_31B and *La*GH13_31B Y295A. (B) Activity of *La*GH13_31 and its Y295A mutant on M3, showing reduced activity of the mutant *La*GH13_31B Y295A as compared to *La*GH13_31B on M3 over time (0, 2, 4, 8, 12, 20, 200 min; S, standard of M2, M3 and M7). (C) Relative fold change in loss of activity of *La*GH13_31B Y295A as compared to *La*GH13_31B on M2 and M3. (D) Transferase kinetics of *La*GH13_31B Y295A on M3 showing the means of triplicates with standard deviations.

## DISCUSSION

Lactobacilli are industrially important and diverse bacteria that colonize a multitude of ecological niches including gastrointestinal tracts of humans and animals. *L. acidophilus* and closely related species from the acidophilus group are adapted to the small intestine of humans, where α-glucans from starch break-down are an abundant metabolic resource (6, 19).

Previously, the maltose phosphorylase *La*GH65 encoded by the MOS gene cluster in *L. acidophilus* NCFM was characterized (20) in addition to a glucan-1,6-α-glucosidase (*La*GH13_31A) from the same strain (21). However, the role of LBA1872, residing in the MOS gene cluster (Fig. 1), remained unclear. The LBA1872 gene product shares amino acid sequence similarities to GH13_31 1,6-α-glucosidases (37% identity to *La*GH13_31A) (21) and to a recently described disproportionating 1,4-α-glucosyltransferase from *Enterococcus fecalis* (MmgT; 35% identity) (25). This tentative functional assignment prompted us to express LBA1872 (*La*GH13_31B) as well as the two other 1,4-α-active enzymes from the MOS gene cluster from *L. acidophilus* NCFM to investigate the roles of these enzymes in MOS metabolism.

### *La*GH13_31B populates a distinct clade in GH13_31, which is defined by a unique loop insertion harbouring a conserved aromatic residue

The valine residue following the catalytic nucleophile in the conserved region II in 1,6-α-glucosidases of GH13_31 (26) is substituted with an alanine in *La*GH13_31B (Fig. 3B, Fig. S6) (21), consistent with the lack of activity of the latter enzyme towards 1,6-α-glucans. Our present phylogenetic tree showed that *La*GH13_31B defines a distinct GH13_31 clade that segregates from characterized GH13_31 members (Fig. 3) and which is populated solely with sequences from lactobacilli. We identified a unique insertion in the loop region between R282 and D305 (*La*GH13_31B numbering), which is lacking in GH13_31 sequences segregating into other clades (Fig. 3). This loop harbours a tyrosine (Y295) that is chemically conserved (Y or F) within the inspected homologues (Fig. S6). The mutational analysis showed that Y295, residing on this loop, is important for activity and substrate affinity based on the loss of curvature of the tranferase activity on the prerred substrate M3 the preferred (Fig. 4D, Fig. S3). The poor or lacking electron density for parts of this loop in *La*GH13_31B are suggestive of its high flexibility. Flexible loops that present substrate binding aromatic residues have been observed in other transferases, e.g. the GH13 4-α-glucanotransferase from *Thermotoga maritima*, which can convert starch, amylopectin and amylose by transferring maltosyl and longer dextrinyl residues to M2 and longer oligosaccharides. This 4-α-glucanotransferase harbors an aromatic “clamp”, which captures substrates at the +1 and +2 subsites (27). In addition, flexible loops that undergo considerable conformational changes during the catalytic cycle have also been identified in 4-α-glucanotransferases of GH77 from *Thermus brockianus* (28), *Escherichia coli* (29) and *Arabidopsis thaliana* (30). In particular, the 4-α-glucanotransferase from *Th. brockianus* was shown to possess an aromatic reside (F251) located on the flexible loop that binds substrates at subsite +1 and +2 (28). The role of the flexible loop in *La*GH13_31B in activity and substrate affinity supports the assignment of this loop as a unique structural and functional motif of the clade defined by *La*GH13_31B, which is conserved within the MOS utilization loci in *Lactobacillus*.

### *La*GH13_31B acts as an enzymatic pivot in the metabolism of maltodextrins in *L. acidophilus* NCFM

The characterization of *La*GH13_31B provides compelling evidence on the 1,4-α-glucosyltransferase activity of the enzyme. The kinetics of disproportionation revealed an exceptionally low activity on M2 and at least a three order of magnitude increase in efficiency (*k*_cat_/*K*_m_) on M3 (Table 1). This difference suggests that the binding of M2 at subsites –1 and +1 is not favorable. By contrast, the roughly 6−9-fold decrease in *K*_m_ when M3 is used an acceptor is consistent with a key role of subsite +2 and the preferred binding mode between subsites –1 through to +2, likely promoted by the additional interactions provided by the aromatic residue (Y295) in the flexible loop discussed above. The preference for M3 is complementary to that of *La*GH13_20, which displays about a 40-fold preference for M4 as compared to M3. Thus, *La*GH13_20 will be more efficient in converting M4 to M2, whereas *La*GH13_31B most likely disproportionates M3 to M2 and to M4. The concerted action of both enzymes will mainly accumulate M2 from odd numbered maltodextrins. The M2 product of these enzymes is phosphorolysed by *La*GH65 to produce glucose (Glc) and Glc-1P, which are further catabolized in glycolysis (Fig. 5). Clearly, *La*GH13_31B contributes to efficient catabolism of odd-numbered maltodextrins through their disproportionation resulting in larger better substrates for the *La*GH13_20 and M2 the preferred substrate for *La*GH65. This mode of catabolism of maltodextrins seems to be widely employed in Firmicutes, but also in other bacteria e.g. *Bifidobacterium*, where a different type of α-glucanotransferase is upregulated during growth on MOS (31). The advantages of having an extra extension step in the breakdown of maltodextrins combined with a GH65 are not obvious, as compared to using α-glucosidases or phosphorylases, which directly break down maltodextrins. One possibility is that the transient extension of maltodextrins, especially during saturation with Glc and lowered rate of *La*GH65 activity due to reverse phosphorolysis (20) may serve as a transient energy reserve, which is more rapidly mobilized than glycogen produced by this organism (32). The accumulation of M2 may also inhibit the activity of the maltogenic α-amylase that is inactive towards this substrate. Further work is needed to verify this role.

**FIG 5.**
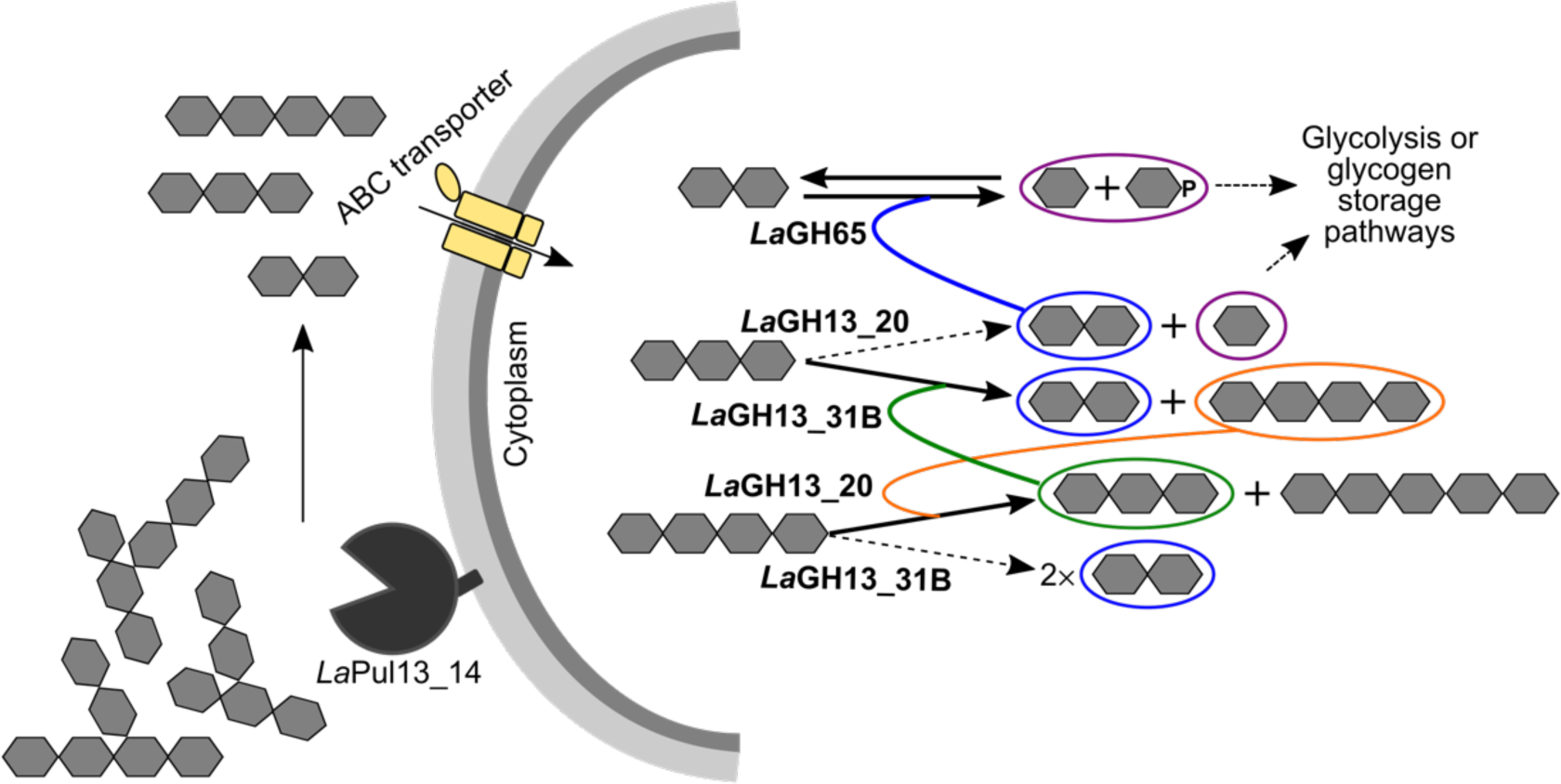
Schematic model of MOS catabolism in *L. acidophilus* NCFM. MOS produced from starch and glycogen degradation by human digestive enzymes, other bacteria or by the extracellular pullulanase (*La*Pul13_14; (19)) are internalised by specific transporters. An ATP-binding cassette transporter is conserved in the locus in *Lactobacillus*, but likely defect in the *L. acidophilus* NCFM due to the presence of a transposase (19). Odd numbered MOS are degraded into M3, whereas even numbered MOS are degraded to M2 by *La*GH13_20. While M3 is a poor substrate for *La*GH13_20, it is preferred by *La*GH13_31B. The action of *La*GH13_31B converts M3 into M2 and M4, which are the preferred substrates for *La*GH65 and *La*GH13_20, respectively. Products are either catabolised via glycolysis (20) or stored as glycogen (32).

## Conclusions

This study provides biochemical data on a previously unknown subfamily of disproportionating enzymes conserved within the maltodextrin metabolism pathway in *Lactobacillus*. Kinetic analysis of the three 1,4-α-active enzymes involved in MOS utilization in *L. acidophilus* revealed that *La*GH13_31B acts as a pivot that may contribute to regulating MOS catabolism by allowing transient storage as longer MOS. We identified a unique signature of this subfamily comprising an insertion in a loop positioned in near proximity to the active site cleft. This loop may act as a clamp that recognizes substrates likely via aromatic stacking at a conserved aromatic side chain.

## MATERIALS AND METHODS

### Chemicals and carbohydrate substrates

High-purity (>95%) chemicals and commercial enzymes were from Sigma-Aldrich (MO, USA) unless otherwise stated. Glucose was purchased from VWR (PA, USA), amylose (DP17) was purchased from Hayashibara Co. (Okayama, Japan).

### Cloning, production and purification of enzymes encoded by the MOS utilization gene cluster in *Lactobacillus acidophilus* NCFM

*L. acidophilus* NCFM genomic DNA, prepared as previously described (20), was used to clone the LBA1871 and LBA1872 genes (GenBank accessions AAV43671.1 and AAV43672.1, respectively) in the pET-21a(+) and pET-28a(+) vectors (Novagen, Darmstadt, Germany), respectively, using primers listed in Table S3 and standard molecular biology protocols. The sequence-verified recombinant vectors designated pET-21a(+)*La*GH13_20 and pET-28a(+)*La*GH13_31B for LBA1871 and LBA1872, respectively, were transformed into *Escherichia coli* production strain Rosetta (DE3) cells (Invitrogen). Production of the maltogenic α-amylase *La*GH13_20 and the oligosaccharide 1,4-α-glucosyltransferase *La*GH13_31B was carried out using a 5 L bioreactor (Biostat B; B. Braun Biotech International, Melsungen, Germany) as previously described (33), with the exception of induction conditions (here *OD*_600_=8, 18°C) and feed (linear gradient 8.4−15 mL h^-1^ in 5 h, and then 15 ml h^-1^ was maintained until harvest). The fermentation was harvested after 48 h of induction (*OD*_600_=50) by centrifugation (15,000*g* for 20 min at 4°C). Portions of 10 g cell pellet were resuspended in 50 mL Bugbuster (BugBuster® Protein Extraction Reagent, Merck Millipore) supplemented with 5 µL Benzonase Nuclease (Novagen) and incubated for 20 min at room temperature. Subsequently the suspension was centrifuged (20,000 *g* for 20 min) and the clarified supernatant was sterile filtered (0.45 µm) and used for purification. Both the *La*GH13_20 and *La*GH13_31B enzymes were purified by immobilized metal ion affinity chromatography on 5 ml HisTrap HP columns (GE Healthcare, Uppsala, Sweden) equilibrated with binding buffer (20 mM HEPES, 500 mM NaCl, 10 mM imidazole, 1 M CaCl_2_, 10% glycerol, pH 7.5) and eluted with a linear gradient of imidazole from 10 mM to 300 mM over 35 CV at a flow rate of 1 mL min^-1^. Fractions enriched with the enzymes were pooled, concentrated (30 kDa Amicon filter; Millipore), and applied to a HiLoad 26/60 Superdex 200 size-exclusion column (GE Healthcare) at a flow rate of 0.5 mL min^-1^. The purifications were performed on an ÄKTA Avant chromatograph (GE Healthcare) and pure enzyme fractions, analyzed by SDS-PAGE, were pooled, concentrated and supplemented with 0.005% NaN_3_ (w/v) after determination of protein concentrations using UV absorbance (*A*_280_, molar extinction coefficient ε_280_=113220 M^-1^ cm^-1^ predicted using the ExPASy server, (34)). *La*GH65 was produced and purified as previously described (20) and further purified using size-exclusion chromatography on a HiLoad 26/60 Superdex 75 column (GE Healthcare). The *La*GH13_31B Y295A variant was generated using primers in Table S3 and the QuickChange II Site-Directed Mutagenesis kit (Agilent) with pET-28a(+)-*La*GH13_31B as template. The mutant was produced in 0.5 L scale using a 2 L baffled shakeflask and purified as described above.

### Differential scanning calorimetry (DSC) stability analysis

The DSC analysis was performed at protein concentrations of 1 mg mL^-1^ in 10 mM sodium phosphate buffer, 150 mM NaCl, pH 6.8, using a Nano DSC (TA instruments). Thermograms were recorded from 20 to 60°C at a scan speed of 1.5°C min^−1^ using buffer as reference. The analysis was done in duplicates. Baseline corrected data were analyzed using the NanoAnalyze software (TA instruments).

### Thin layer chromatography (TLC) of enzyme product profiles

Disproportionating abilities of *La*GH13_31B (0.5 µM) and *La*GH13_31B Y295A (0.5 µM) on M3 (10 mM) were visualized by TLC. Reactions (100 µL) were incubated in standard assay buffer at 37°C and 2 µL aliquots were removed at appropriate time points and spotted on a silica gel 60 F454 plate (Merck). The separation was carried out in butanol:ethanol:milliQ water (5:3:2) (v/v) as mobile phase and sugars were visualized with 5-methylresorcinol:ethanol:sulfuric acid (2:80:10) (v/v) and heat treatment.

### Enzyme activity

Oligo-1,6-*α*-glucosidase activity by *La*GH13_31B (50 nM) was analyzed at pH 6.8 and 37°C for 12 min in 300 µL reactions containing isomaltose or panose (5 mM) in the assay buffer (10 mM MES, 150 mM NaCl, 0.005% Tween20, pH 6.8). The liberated Glc was quantified using a modified glucose oxidase/peroxidase assay (GOPOD; Megazyme, Wicklow, Ireland) (21). Similarly, maltose phosphorolysis kinetics by *La*GH65 was analyzed with reactions containing M2 (0.63−20 mM) and *La*GH65 (23 nM) and the assay buffer (100 mM phosphate/citrate, 0.005% Tween20). Aliquots (50 µL) were removed at five time points (3, 6, 9, 12, and 15 min) and added to 100 µL 2 M Tris-HCl, pH 7, to stop the reaction, and Glc was quantified as described above. The same assay was used to measure the hydrolysis kinetics of *La*GH13_20 on M3 (0.63−20 mM) and the disproportionation activity of *La*GH13_31B on M2 (1.6−300 mM), with the only exception that *La*GH13_20 (70 nM) and *La*GH13_31B (6 µM) were incubated in the assay buffer (10 mM MES, 150 mM NaCl, 0.005% Tween20, pH 6.8). The hydrolysis kinetics of *La*GH13_20 (7 nM) towards M4 (0.63−20 mM) and disproportionation kinetics of *La*GH13_31B (3.8 nM) on M3 (0.63−30 mM) were determined using a coupled assay, where 0.2 µM *La*GH65 and 20 mM phosphate were included in the assay buffer.

A similar assay was also applied to measure the disproportionation kinetics of M3 (0.63−40 mM) and M4 (1.3−40 mM) with 1.9 and 3.8 nM of *La*GH13_31B, respectively, using HPAEC-PAD as described below. Appropriate amounts of reaction mixtures were diluted into 0.1 M NaOH before injection and the separated saccharides were quantified based on peak areas. Glc, M2, and MOS (0.015–1 mM) were used as standards. The Michaelis-Menten model was fit to the initial rate data to derive kinetic parameters using OriginPro 2015 software (OriginLab, Northampton, MA). The utilization of MOS by *La*GH13_31B (3.8 nM) alone, or by a mixture of *La*GH65 (3.8 nM), *La*GH13_20 (3.8 nM) and *La*GH13_31B (3.8 nM) towards 5 mM of either M3, M4 or M5 were also analyzed by HPAEC-PAD. Similar assay conditions as described above was applied, with the only exception that samples were incubated for 24 h in the buffer of the coupled assay.

Relative disproportionation activities of *La*GH13_31 (60 µM) and *La*GH13_31 Y295A (114 µM) on M2 (100 mM) were analyzed as described above, using standard assay buffer (10 mM MES, 150 mM NaCl, 0.005% Tween20, pH 6.8) and an incubation time of 10 min at 37°C. The liberated Glc was quantified using the coupled enzyme assay described above. The analysis was performed in technical triplicates. Relative activities of *La*GH13_31 (5 nM) and *La*GH13_31 Y295A (0.5 µM) towards M3 (5 mM) were assayed under similar reaction conditions. For determining initial rates, reactions (150 µL) were incubated for 8 min at 37°C and aliquots of 15 µL were removed every minute and quenched in 135 µL 0.1 M NaOH. Reaction products were quantified using HPAEC-PAD as in detail described below. A similar assay was used to measure the disproportionation kinetics of *La*GH13_31 Y295A on M3 (1−60 mM), with the only exception that a single aliquot was removed after 5 min of incubation before quenching in 0.1 M NaOH. The HPAEC-PAD analysis was performed in technical triplicates.

### High pressure anion exchange chromatography with pulsed amperometric detection (HPAEC-PAD)

1,6-α-glucosidase activity and enzyme kinetics were analyzed using HPAEC-PAD analysis. Samples (10 µL) were injected into a Carbopac PA200 analytical column coupled with a guard column (Thermo Fisher Scientific, Sunnyvale, USA) installed on an ICS-3000 chromatograph (Thermo Fisher Scientific) and analyzed at a flow rate of 0.35 mL min^-1^. The elution was carried out using a constant concentration of 100 mM NaOH and, in addition, from 0–10 min a gradient of 40–150 mM sodium acetate (NaOAc); 10–11 min, 150–400 mM NaOAc; 11–15 min, 400 mM NaOAc; 15–20 min, a linear gradient from 400 mM to initial conditions of 40 mM NaOAc. The system was interfaced using a Chromeleon version 6.7, which was also used to evaluate the chromatograms.

### Sequence alignment and phylogenetic analysis

All lactobacilli protein sequences classified into GH13_31 in the CAZy database (12) together with the sequences of the GH13_31 members classified as characterized were retrieved from CAZy database. Redundancy of the protein sequences was reduced using the Decrease redundancy server (web.expasy.org/decrease_redundancy/) using 99% max identity as a rule to reduce redundancy. Then a structure-guided multiple sequence alignment was made using the PROMALS3D webserver (35) including available structures of GH13_31 enzymes using default settings. Based on the structure-based multiple sequence alignment a phylogenetic tree was constructed using the Maximum Likelihood based on conserved sites using the Jones, Taylor, and Thorton (JTT) model and with 500 bootstrap replications. The tree construction was done using MEGA X (36) and visualized using Dendroscope version 3.6.3 (37). The multiple sequence alignment was visualized using ESPript 3.0 (38).

The organization of the MOS gene cluster of different *Lactobacillus* strains was analyzed using the genome database provided by the National Center for Biotechnology Information (NCBI), MGcV: the microbial genomic context viewer for comparative genome analysis (39), KEGG: Kyoto Encyclopedia of Genes and Genomes (40) and previous studies (20, 41).

### Crystallization, data collection and structure determination

Screening for crystallization conditions was performed using 3.6 mg mL^-1^ *La*GH13_31B in 10 mM MES, pH 6.5, 10 mM NaCl, and 0.5 mM CaCl_2_ by the sitting-drop vapour-diffusion method using the JCSG+ (Qiagen, Hilden, Germany), Index (Hampton Research, CA US), and Morpheus® MD1–46 (Molecular Dimensions, Newmarket, UK) screens. An Oryx8 liquid-handling robot (Douglas Instruments, Hungerford, UK) was used to set up the screens in MRC 2-drop plates (Douglas Instruments, Hungerford, UK) with a total drop volume of 0.3 µL and 3:1 and 1:1 ratios of protein solution and reservoir solution at room temperature. Small thin needle crystals appeared within two weeks at room temperature with a reservoir containing an equimolar mixture (0.02 M each) of sodium L-glutamate, DL-alanine, glycine, DL-lysine HCl, DL-serine), buffer system 3 pH 8.5 (0.1 M Bicine and 0.1 M Trizma base), 20% v/v PEG 500 MME; 10% w/v PEG 20000, pH 8.5 (Morpheus®) in a protein:reservoir (1:1) droplet. Microseed matrix screening using the above needles (42) was performed in several of the above screens, but suitable crystals only appeared under the same conditions as above in several of the above screens. No extra cryoprotection was used before the crystals were mounted and flash frozen in liquid nitrogen. Diffraction data were collected at the ESRF beamline ID23-1 and processed with the XDS package (43) in space group *P*2_1_ to 2.8 Å, with cell and processing statistics as reported in Table 2. Molecular replacement was carried out in Molrep (44) with the structure of a GH13_31 oligo-1,6-*α*-glucosidase (PDB: 1UOK) as model, and 2 molecules/asymmetric unit were identified as suggested by Matthew’s number. The electron density generated from the solution showed a Ca^2+^ binding site, absent in the search model, but present in several homologues. Several areas, especially in loop regions, had initially extremely poor density. The structure was refined using both Phenix (45) and REFMAC 5.0 (46) and with the aid of average maps from COOT (47) especially at the initial stages of model building and refinement, whereas several autobuild strategies in phenix, jelly-body, and ProSmart restrained refinement (using GH13_31 structures with PDB: 4AIE and 4MB1) in REFMAC 5.0 were applied for the last stages. Non-crystallographic symmetry (NCS) restrains have been used for most of the structure. The loop region 286–295 (comprising part of the unique loop containing Y295) could not be modelled confidently though some residual density is clearly seen in this region (Fig. S5). No density was visible for loop 519–522. Other regions have poor density in one of the two copies in the asymmetric unit, and here the area has been modelled similarly to the corresponding regions in the other chain, in which the density is considerably better. Structures were visualized with PyMOL, version 2.1.1 (Schrödinger, LLC). Atomic coordinates of *La*GH13_31B have been deposited at the Protein Data Bank (accession: 6Y9T).

## SUPPLEMENTAL MATERIAL

Supplemental material for this article may be found at the AEM website.

## ACKNOWLEDGEMENTS

Mette Pries (Technical University of Denmark) and Dorte Boelskifte (University of Copenhagen) are thanked for technical assistance. This work was supported by a PhD scholarship from the Technical University of Denmark to SA and a FøSu grant from the Danish Strategic Research Council to the project “*Gene discovery and molecular interactions in prebiotics/probiotics systems. Focus on carbohydrate prebiotics*”. The Carlsberg Foundation is acknowledged for an instrument grant that funded the DSC equipment. The Danish Ministry of Higher Education and Science through the Instrument Center DANSCATT funded travel to synchrotrons. The crystallographic experiments were performed on beamline ID23-1 at the European Synchrotron Radiation Facility (ESRF), Grenoble, France. We are grateful to the ESRF staff for assistance.

## REFERENCES

1. Muegge BD, Kuczynski J, Knights D, Clemente JC, Gonzalez A, Fontana L, Henrissat B, Knight R, Gordon JI. 2011. Diet drives convergence in gut microbiome functions across mammalian phylogeny and within humans. Science 332:970–974.

2. Thaiss CA, Zmora N, Levy M, Elinav E. 2016. The microbiome and innate immunity. Nature 535:65–74.

3. Spanogiannopoulos P, Bess EN, Carmody RN, Turnbaugh PJ. 2016. The microbial pharmacists within us: A metagenomic view of xenobiotic metabolism. Nat Rev Microbiol 14:273–287.

4. Flint HJ, Bayer EA, Rincon MT, Lamed R, White BA. 2008. Polysaccharide utilization by gut bacteria: potential for new insights from genomic analysis. Nat Rev Microbiol 6:121–31.

5. David LA, Maurice CF, Carmody RN, Gootenberg DB, Button JE, Wolfe BE, Ling A V, Devlin AS, Varma Y, Fischbach MA, Biddinger SB, Dutton RJ, Turnbaugh PJ. 2014. Diet rapidly and reproducibly alters the human gut microbiome. Nature 505:559–563.

6. Donaldson GP, Lee SM, Mazmanian SK. 2016. Gut biogeography of the bacterial microbiota. Nat Rev Microbiol 14:20–32.

7. Sanders ME, Klaenhammer TR. 2001. Invited review: the scientific basis of *Lactobacillus acidophilus* NCFM functionality as a probiotic. J Dairy Sci 84:319–31.

8. Lyra A, Hillilä M, Huttunen T, Männikkö S, Taalikka M, Tennilä J, Tarpila A, Lahtinen S, Ouwehand AC, Veijola L. 2016. Irritable bowel syndrome symptom severity improves equally with probiotic and placebo. World J Gastroenterol 22:10631–10642.

9. Altermann E, Russell WM, Azcarate-Peril MA, Barrangou R, Buck BL, McAuliffe O, Souther N, Dobson A, Duong T, Callanan M, Lick S, Hamrick A, Cano R, Klaenhammer TR. 2005. Complete genome sequence of the probiotic lactic acid bacterium *Lactobacillus acidophilus* NCFM. Proc Natl Acad Sci U S A 102:3906–3912.

10. Ouwehand AC, Tiihonen K, Saarinen M, Putaala H, Rautonen N. 2009. Influence of a combination of *Lactobacillus acidophilus* NCFM and lactitol on healthy elderly: Intestinal and immune parameters. Br J Nutr 101:367–375.

11. Magro DO, De Oliveira LMR, Bernasconi I, Ruela MDS, Credidio L, Barcelos IK, Leal RF, Ayrizono MDLS, Fagundes JJ, Teixeira LDB, Ouwehand AC, Coy CSR. 2014. Effect of yogurt containing polydextrose, *Lactobacillus acidophilus* NCFM and *Bifidobacterium lactis* HN019: A randomized, double-blind, controlled study in chronic constipation. Nutr J 13:1–5.

12. Lombard V, Ramulu HG, Drula E, Coutinho PM, Henrissat B. 2014. The carbohydrate-active enzymes database (CAZy) in 2013. Nucleic Acids Res 42:490–495.

13. Andersen JM, Barrangou R, Abou Hachem M, Lahtinen S, Goh YJ, Svensson B, Klaenhammer TR. 2011. Transcriptional and functional analysis of galactooligosaccharide uptake by *lacS* in *Lactobacillus acidophilus*. Proc Natl Acad Sci 108:17785–17790.

14. Andersen JM, Barrangou R, Abou Hachem M, Lahtinen SJ, Goh Y-J, Svensson B, Klaenhammer TR. 2012. Transcriptional analysis of prebiotic uptake and catabolism by *Lactobacillus acidophilus* NCFM. PLoS One 7:e44409.

15. Theilmann MC, Goh YJ, Nielsen KF, Klaenhammer TR, Barrangou R, Abou Hachem M. 2017. *Lactobacillus acidophilus* metabolizes dietary plant glucosides and externalizes their bioactive phytochemicals. MBio 8:1–15.

16. Bertoft E. 2017. Understanding starch structure: Recent progress. Agronomy 7:56.

17. Ze X, Duncan SH, Louis P, Flint HJ. 2012. *Ruminococcus bromii* is a keystone species for the degradation of resistant starch in the human colon. ISME J 6:1535–1543.

18. Cockburn DW, Orlovsky NI, Foley MH, Kwiatkowski KJ, Bahr CM, Maynard M, Demeler B, Koropatkin NM. 2015. Molecular details of a starch utilization pathway in the human gut symbiont *Eubacterium rectale*. Mol Microbiol 95:209–230.

19. Møller MS, Goh YJ, Rasmussen KB, Cypryk W, Celebioglu HU, Klaenhammer TR, Svensson B, Abou Hachem M. 2017. An extracellular cell-attached pullulanase confers branched α-glucan utilization in human gut *Lactobacillus acidophilus*. Appl Environ Microbiol 83:1–13.

20. Nakai H, Baumann MJ, Petersen BO, Westphal Y, Schols H, Dilokpimol A, Abou Hachem M, Lahtinen SJ, Duus JØ, Svensson B. 2009. The maltodextrin transport system and metabolism in *Lactobacillus acidophilus* NCFM and production of novel α-glucosides through reverse phosphorolysis by maltose phosphorylase. FEBS J 276:7353–7365.

21. Møller MS, Fredslund F, Majumder A, Nakai H, Poulsen J-CN, Lo Leggio L, Svensson B, Abou Hachem M. 2012. Enzymology and structure of the GH13_31 glucan 1,6-α-glucosidase that confers isomaltooligosaccharide utilization in the probiotic *Lactobacillus acidophilus* NCFM. J Bacteriol 194:4249–4259.

22. Auiewiriyanukul W, Saburi W, Kato K, Yao M, Mori H. 2018. Function and structure of GH13_31 α-glucosidase with high α-(1→4)-glucosidic linkage specificity and transglucosylation activity. FEBS Lett 592:2268–2281.

23. Shirai T, Hung VS, Morinaka K, Kobayashi T, Ito S. 2008. Crystal structure of GH13 α-glucosidase GSJ from one of the deepest sea bacteria. Proteins 73:126–33.

24. Oslancová A, Janecek S. 2002. Oligo-1,6-glucosidase and neopullulanase enzyme subfamilies from the α-amylase family defined by the fifth conserved sequence region. Cell Mol Life Sci 59:1945–59.

25. Joyet P, Mokhtari A, Riboulet-Bisson E, Blancato VS, Espariz M, Magni C, Hartke A, Deutscher J, Sauvageot N. 2017. Enzymes required for maltodextrin catabolism in *Enterococcus faecalis* exhibit novel activities. Appl Environ Microbiol 83:1–15.

26. Janeček Š. 2002. How many conserved sequence regions are there in the α-amylase family? Biologia (Bratisl) 57:29–41.

27. Roujeinikova A, Raasch C, Sedelnikova S, Liebl W, Rice DW. 2002. Crystal structure of *Thermotoga maritima* 4-α-glucanotransferase and its acarbose complex: implications for substrate specificity and catalysis. J Mol Biol 321:149–162.

28. Jung J-H, Jung T-Y, Seo D-H, Yoon S-M, Choi H-C, Park BC, Park C-S, Woo E-J. 2011. Structural and functional analysis of substrate recognition by the 250s loop in amylomaltase from *Thermus brockianus*. Proteins 79:633–44.

29. Weiss SC, Skerra A, Schiefner A. 2015. Structural basis for the interconversion of maltodextrins by MalQ, the amylomaltase of *Escherichia coli*. J Biol Chem 290:21352–21364.

30. O’Neill EC, Stevenson CEM, Tantanarat K, Latousakis D, Donaldson MI, Rejzek M, Nepogodiev SA, Limpaseni T, Field RA, Lawson DM. 2015. Structural dissection of the maltodextrin disproportionation cycle of the *Arabidopsis* plastidial enzyme DPE1. J Biol Chem 290:29834–29853.

31. Andersen JM, Barrangou R, Abou Hachem M, Lahtinen SJ, Goh YJ, Svensson B, Klaenhammer TR. 2013. Transcriptional analysis of oligosaccharide utilization by *Bifidobacterium lactis* Bl-04. BMC Genomics 14:312.

32. Goh YJ, Klaenhammer TR. 2014. Insights into glycogen metabolism in *Lactobacillus acidophilus*: impact on carbohydrate metabolism, stress tolerance and gut retention. Microb Cell Fact 13:94.

33. Fredslund F, Abou Hachem M, Jonsgaard Larsen R, Gerd Sørensen P, Coutinho PM, Lo Leggio L, Svensson B. 2011. Crystal structure of α-galactosidase from *Lactobacillus acidophilus* NCFM: Insight into tetramer formation and substrate binding. J Mol Biol 412:466–480.

34. Gasteiger E, Gattiker A, Hoogland C, Ivanyi I, Appel RD, Bairoch A. 2003. ExPASy: The proteomics server for in-depth protein knowledge and analysis. Nucleic Acids Res 31:3784–3788.

35. Pei J, Kim B-H, Grishin N V. 2008. PROMALS3D: a tool for multiple protein sequence and structure alignments. Nucleic Acids Res 36:2295–300.

36. Kumar S, Stecher G, Li M, Knyaz C, Tamura K. 2018. MEGA X: Molecular evolutionary genetics analysis across computing platforms. Mol Biol Evol 35:1547–1549.

37. Huson DH, Scornavacca C. 2012. Dendroscope 3: An interactive tool for rooted phylogenetic trees and networks. Syst Biol 61:1061–1067.

38. Robert X, Gouet P. 2014. Deciphering key features in protein structures with the new ENDscript server. Nucleic Acids Res 42:W320–W324.

39. Overmars L, Kerkhoven R, Siezen RJ, Francke C. 2013. MGcV: the microbial genomic context viewer for comparative genome analysis. BMC Genomics 14:209.

40. Kanehisa M, Goto S, Hattori M, Aoki-Kinoshita KF, Itho M, Kawashima S, Katayama T, Araki M, Hirakawa M. 2006. From genomics to chemical genomics: new developments in KEGG. Nucleic Acids Res 34:D354–D357.

41. Monedero V, Yebra MJ, Poncet S, Deutscher J. 2008. Maltose transport in *Lactobacillus casei* and its regulation by inducer exclusion. Res Microbiol 159:94–102.

42. D’Arcy A, Bergfors T, Cowan-Jacob SW, Marsh M. 2014. Microseed matrix screening for optimization in protein crystallization: What have we learned? Acta Crystallogr Sect Struct Biol Commun 70:1117–1126.

43. Kabsch W. 2010. XDS. Acta Crystallogr D Biol Crystallogr 66:125–132.

44. Vagin A, Teplyakov A. 2010. Molecular replacement with MOLREP. Acta Crystallogr Sect D Biol Crystallogr 66:22–25.

45. Adams PD, Afonine P V, Bunkóczi G, Chen VB, Davis IW, Echols N, Headd JJ, Hung L-W, Kapral GJ, Grosse-Kunstleve RW, McCoy AJ, Moriarty NW, Oeffner R, Read RJ, Richardson DC, Richardson JS, Terwilliger TC, Zwart PH. 2010. PHENIX: a comprehensive Python-based system for macromolecular structure solution. Acta Crystallogr D Biol Crystallogr 66:213–221.

46. Nicholls RA, Long F, Murshudov GN. 2013. Recent advances in low resolution refinement tools in REFMAC5, p. 231–258. In Read, R, Urzhumtsev, AG, Lunin, VY (eds.), Advancing Methods for Biomolecular Crystallography. Springer Netherlands, Dordrecht.

47. Emsley P, Lohkamp B, Scott WG, Cowtan K. 2010. Features and development of Coot. Acta Crystallogr D Biol Crystallogr 66:486–501.

48. MacGregor EA, Janecek S, Svensson B. 2001. Relationship of sequence and structure to specificity in the α-amylase family of enzymes. Biochim Biophys Acta 1546:1–20.

